# N-glycosylation network construction and analysis to modify glycans on the spike S glycoprotein of SARS-CoV-2

**DOI:** 10.1101/2020.06.23.167791

**Authors:** Sridevi Krishnan, Giri P Krishnan

## Abstract

**Background:** The spike S-protein of SARS-CoV-2 is N-glycosylated. The N-glycan structure and composition of this glycoprotein influence how the virus interacts with host cells.

**Objective:** To identify a putative N-glycan biosynthesis pathway of SARS-CoV-2 (HEK293 cell recombinant) from previously published mass spectrometric studies, and to identify what effect blocking some enzymes has on the overall glycoprotein profile. Finally, our goal was to provide the biosynthesis network, and glycans in easy-to-use format for further glycoinformatics work.

**Methods:** We reconstructed the glycosylation network based on previously published empirical data using GNAT, a glycosylation network analysis tool. Our compilation of the network tool had 23 glycosyltransferase and glucosidase enzymes, and could infer the pathway of glycosylation machinery based on glycans identified in the virus spike protein. Once the glycan biosynthesis pathway was generated, we simulated the effect of blocking specific enzymes - Mannosidase-II and alpha-1,6-fucosyltransferase to see how they would affect the biosynthesis network.

**Results:** Of the 23 enzymes, a total of 12 were involved in glycosylation of SARS-CoV-2 - Man-Ia, MGAT1, MGAT2, MGAT4, MGAT5, B4GalT, B4GalT, Man II, SiaT, ST3GalI, ST3GalVI and FucT8. Blocking enzymes resulted in a substantially modified glycan profile of the protein.

**Conclusions:** A network analysis of N-glycan biosynthesis of SARS-CoV-2 spike protein shows an elaborate enzymatic pathway with several intermediate glycans, along with the ones identified by mass spectrometric studies. Variations in the final N-glycan profile of the virus, given its site-specific microheterogeneity, could be a factor in the host response to the infection and response to antibodies. Here we provide all the resources generated - the glycans derived from mass spectrometry and intermediate glycans in glycoCT xml format, and the biosynthesis network for future drug and vaccine development work.

## Introduction

Glycosylation is a very common and complex post-translational modification process of proteins^1^. About 90% of human proteins are N-glycosylated. N-glycosylation has been shown to have an impact on protein folding and function in some cases^2^, but not in others^3^. Among various reasons, this could suggest that the site of glycosylation, or the type of the glycan – mannose-rich, hybrid or complex, also play a role in influencing protein structure and function. Viruses use the host glycosylation machinery to replicate. So, it can be useful to understand which enzymes are critical to this process. Bioinformatics tools have been used to determine physiological processes in the field of genetics^4^ and cancer biology^5^ with success, and glycobiology is no exception^6^.

Given the current SARS-CoV-2 pandemic and the role of the spike-S-glycoprotein in the virus entry and infection of host cells^7^, we chose to map the enzymatic machinery that is responsible for the spike-S-glycoprotein synthesis. This pathway could help with vaccine development, where modified glycoproteins can be used to simulate binding using molecular dynamic binding studies with immune cells or immunoproteins, determining the host response^8^. Several groups of researchers have developed bioinformatics tools that are useful in understanding glycosylation, both for glycosite prediction^9^ and to predict the bio-synthetic pathway based on empirical data from mass spectrometry^10^. GNAT^11^, which is used in the current manuscript, is a tool that allows to selectively identify glycosylation pathways using the glycan profile of each protein based on specific enzyme rules and constraints.

The glycosylation profile of the spike S protein has been reported by three independent research groups thus far, using sequencing, mass spectrometry and imaging tools^12–14^. There is some variability in the reported glycosylation profile based on these reports. And further research is necessary to find out how the host processes are actually being used to generate the spike glycoprotein, as pointed out by Shajahan et al^14^.

In this work, we examined the effect of blocking two different glycosylation enzymes, to see if we can modify the network of glycans developed as part of the virus spike glycoprotein. Specifically, we chose to simulate blocking Mannosidase-II and alpha-1,6-fucosyltransferase. Expanding on these blocking studies can help identify the ideal targets to choose that affect the virus replication, since that is dependent on the spike glycoprotein, without affecting the host.

## Methods

### Simulated N-glycan biosynthetic network generation

We used the list of the most representative N-glycans per glycosite detected by Zhang et al (obtained from figure 3 in their manuscript)^12^, and the N-glycan profile of SARS-CoV-2 as reported by Shahjahan et al (obtained from figure 4 in their manuscript)^14^ to generate our primary glycosylation biosynthesis networks. Data from Zhang et al^11^ and Shajahan et al^13^ were both obtained from recombinant viral proteins expressed in HEK293 cells. We generated glycoCT xml version of these glycans using the glycanbuilder tool^15^ and verified structures using the glycan chemNIST MS database^16^. Our supplemental documents provide all the glycans that were used and generated as part of this analysis in glycoCT xml format.

### GNAT and Inferglycan pathway

As mentioned earlier, we used GNAT^11^, with the additional enzymes developed by Hou et al^17^ to have a functional simulation tool for N-glycan biosynthesis with a total of 23 enzymes. The reverse inference algorithm derives the N-glycosylation pathway from a given set of reaction products and possible enzymes. For each enzyme, a set of rules and constraints is provided that defines its action. Given a glycan, it is then possible to identify a set of reactions that led to the production of the glycan, by examining all the enzyme rules with constraints in reverse. This will generate either a network of glycans which are predecessors of the given glycan all the way to Man-9, which is the ‘parent’ glycan for this pathway, or will result in an empty set, i.e., the given glycan has no predecessor leading the way to Man-9. Since we provided all the required enzymes to construct the entire glycosylation pathway, we were able to identify all the paths from Man-9 to all the glycans observed in the Mass Spectrometry studies. Since, there is no good way to determine the linkage information (i.e. structure) from mass spectrometry-based composition^18^, there is likely structural heterogeneity that we failed to account for. This heterogeneity can also affect the pathway that was chosen or the enzyme that was involved in the biosynthesis, which is a limitation in our current approach.

### Simulated inhibition of glycan biosynthesis pathway

There are established chemical inhibitors for blocking or slowing down the rate of glycosylation biosynthetic reactions^19^. The blocking of these pathways has been discussed and considered in the past^20,21^. We identified the N-glycan biosynthesis pathway using GNAT, based on which we chose to simulate blocking of ManII and FucT8. We generated network graphs with glycans as nodes, and the enzymes as edges with and without the simulated enzyme inhibition.

## Results

**Figure 1** depicts the network of reactions (edges colored by enzyme) and glycans (nodes, numbered ones were identified by the masspec data) generated based on glycans from Zhang et al. The glycans detected by mass spectrometric analyses are numbered, and the intermediate ones generated in the biosynthesis pathway are not numbered. Our results suggest the involvement of the following 9 enzymes: Man-Ia, Man II, MGAT2, MGAT3, MGAT4, MGAT5, B4GalT, SiaT and FucT8. In addition to the full network of biosynthetic reactions, Figure 2 also has panels depicting the effect blocking Man-II and FucT8. In both cases, only 3 of the total 10 glycans were formed, thus modifying the glycan profile of the viral protein. In panel B and C, the network depicting 3 glycans (1, 4 and 9) that result in being traced back to the Man-9 residue (residue # 102) will be formed. The other network that is independent of this (not containing resiude#102) comprises all the glycans that would not be formed. **Table 1** presents the glycans that could and could not be formed when Man II and FucT8 were blocked. As previously mentioned, these two inhibitors were chosen because they have been established to specifically inhibit glycosylation enzymes.

**Figure 1:**
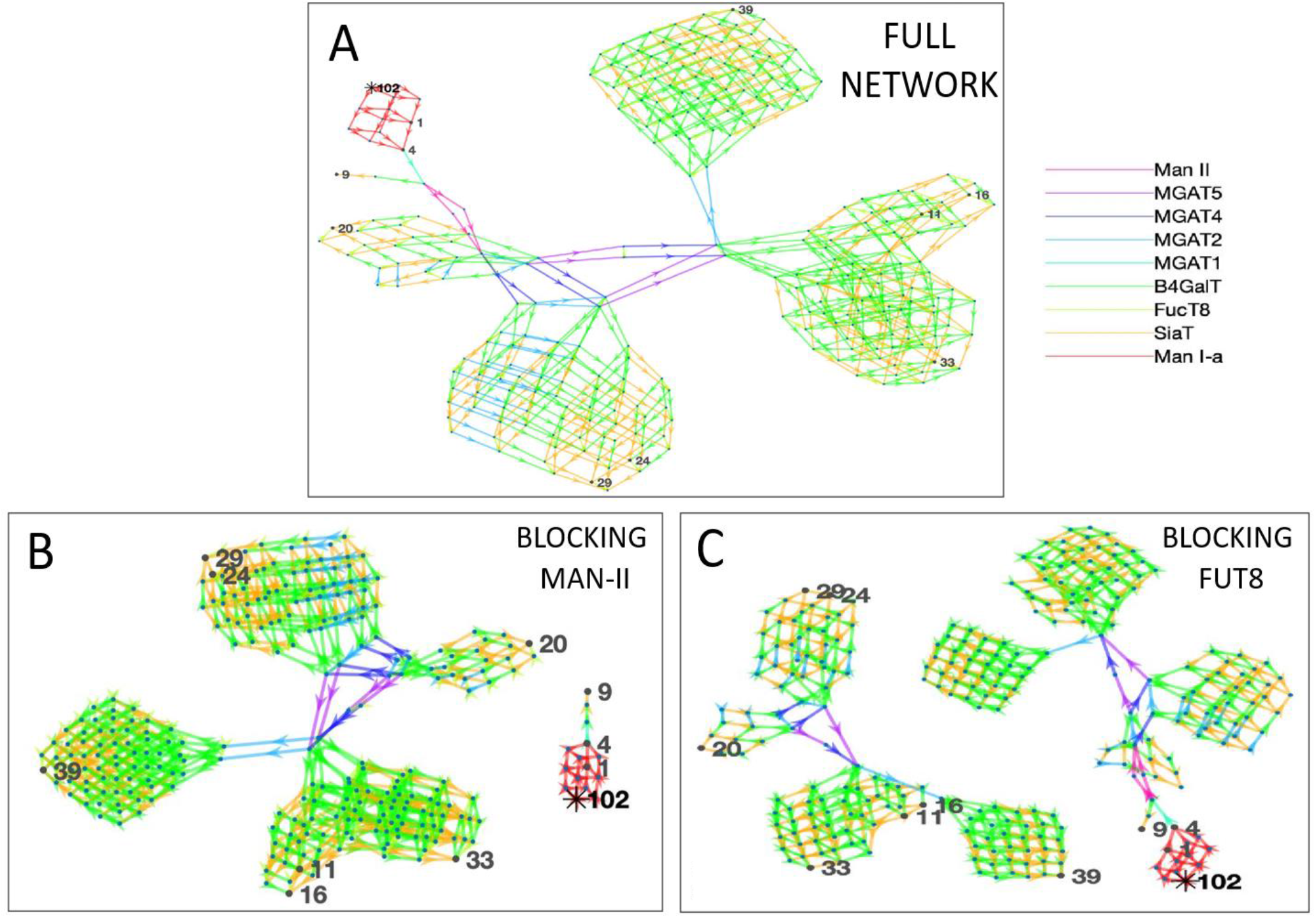
The network of glycans along with the enzymes involved in generating the 10 most abundant glycans as per Zhang et al (panel A). All the 10 glycans (identified by nodes 1, 4, 9, 11, 16, 20, 24, 29, 33 and 39) were traced back to their Man-9 parent glycan (node 102) and the biosynthesis pathway involved the use of 9 enzymes as presented in the key to the right, as denoted by the colors. Panels B and C display the disruption of the biosynthesis network when simulating blocking of alpha-mannosidase (Man-II -Panel B) or alpha-1,6-fucosyltransferase (FucT8-Panel C). In the network generated with enzymes in panel B and C, only 3 of the total 10 glycans are formed.

**Figure 2:**
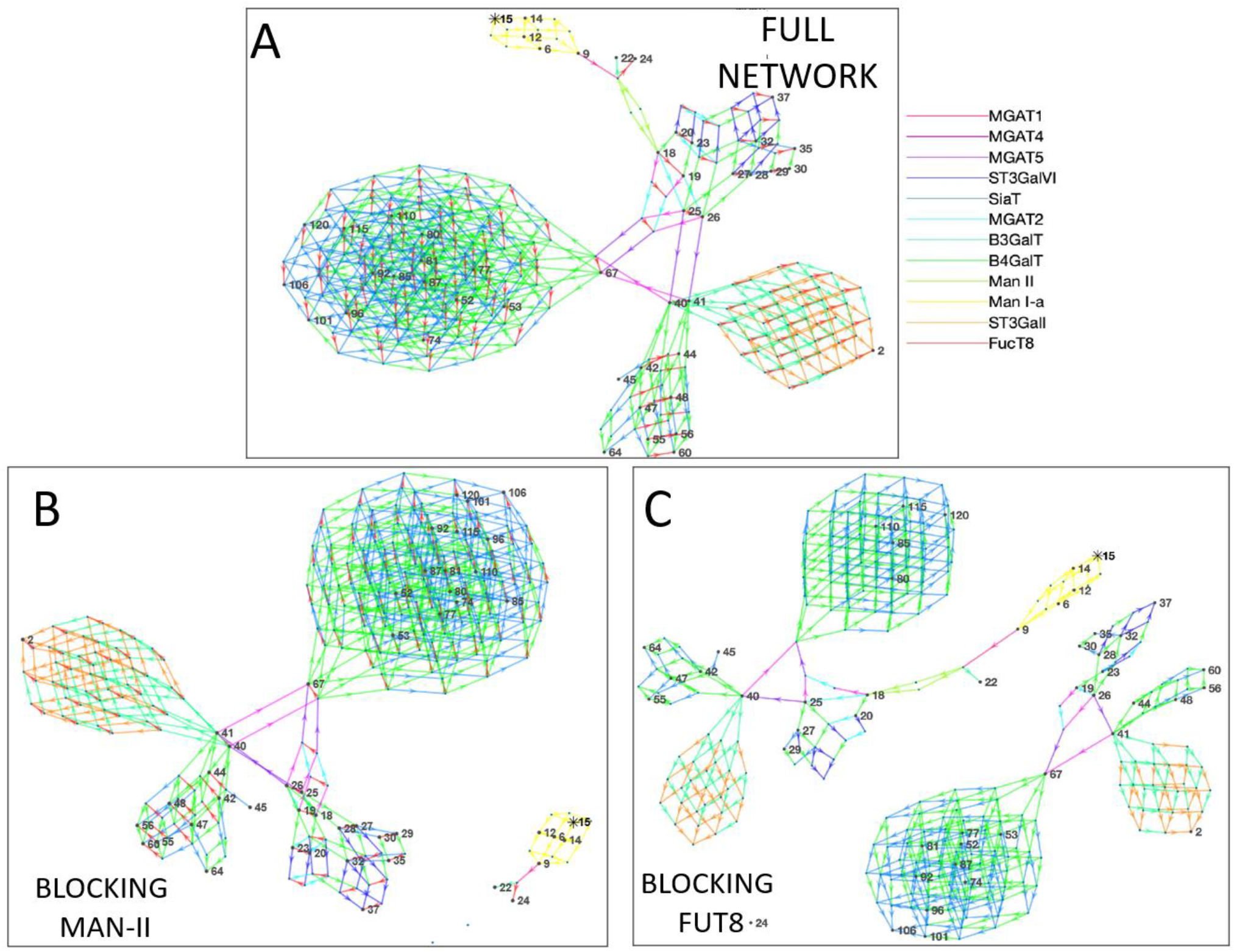
The network of glycans along with the enzymes involved in generating the glycans as per Shajahan et al (Panel A). All the 48 glycans (identified by numbered nodes) were traced back to their Man-9 parent glycan (node 15). The colors (explained in the key) depict the different enzymes involved in the biosynthesis pathway. A total of 12 N-glycan biosynthesis enzymes were required to develop this profile of glycans. Panels B and C display disruption of the network of glycans generated when simulating blocking alpha-mannosidase (Man-II - Panel B) or alpha-1,6-fucosyltransferase (FucT8 -Panel C). In the network generated with enzymes in panel B only 5 glycans are formed, and in panel C only 21 of the 48 are formed.

**Table 1:** List of glycans that were formed vs not formed when specific enzymes were blocked when using data from Zhang et al. Glycan numbers indicate nodes in Figure 1.

| **Blocking FucT8** |  |
| --- | --- |
| **Formed** | **Not-Formed** |
| Glycan: 4 HexNAc2Hex5 | Glycan :11 HexNac6hex6fuc1sia1 |
| Glycan: 1 HexNAc2Hex6 | Glycan :16 HexNac6hex6fuc1sia2 |
| Glycan: 9 HexNAc3Hex6NeuAc1 | Glycan :20 HexNAc4Hex5Fuc1NeuAc2 |
|  | Glycan :24 HexNAc5Hex6Fuc1NeuAc2 |
|  | Glycan :29 HexNAc5Hex6Fuc1NeuAc3 |
|  | Glycan :33 HexNAc6Hex7Fuc1NeuAc2 |
|  | Glycan :34 HexNAc7Hex7Fuc1NeuAc2 |
| **Blocking ManII** |  |
| **Formed** | **Not-Formed** |
| Glycan :4 HexNAc2Hex5 | Glycan :11 HexNac6hex6fuc1sia1 |
| Glycan :1 HexNAc2Hex6 | Glycan :16 HexNac6hex6fuc1sia2 |
| Glycan :9 HexNAc3Hex6NeuAc1 | Glycan :20 HexNAc4Hex5Fuc1NeuAc2 |
|  | Glycan :24 HexNAc5Hex6Fuc1NeuAc2 |
|  | Glycan :29 HexNAc5Hex6Fuc1NeuAc3 |
|  | Glycan :33 HexNAc6Hex7Fuc1NeuAc2 |
|  | Glycan :39 HexNAc7Hex7Fuc1NeuAc2 |

**Figure 2** depicts the network of reactions and glycans similar to Figure 2, but is based on glycans from Shajahan et al. Similar to Figure 1, the glycans detected by mass spectrometric analyses are numbered, and the intermediate ones generated in the biosynthesis pathway are not. Here, 12 enzymes were involved – Man-Ia, MGAT1, MGAT2, MGAT4, MGAT5, B4GalT, B4GalT, Man II, SiaT, ST3GalI, ST3GalVI and FucT8. Similar to Figure 2, blocking of Man II and FucT8 resulted in a very different glycosylation profile of the overall protein, since only the network of glycans that can be traced back to Man-9 (residue #15) would be formed, and the others (networks without residue #15) would not be formed. **Table 2** presents the glycans that could and could not be formed when Man II and FucT8 were blocked.

**Table 2:**
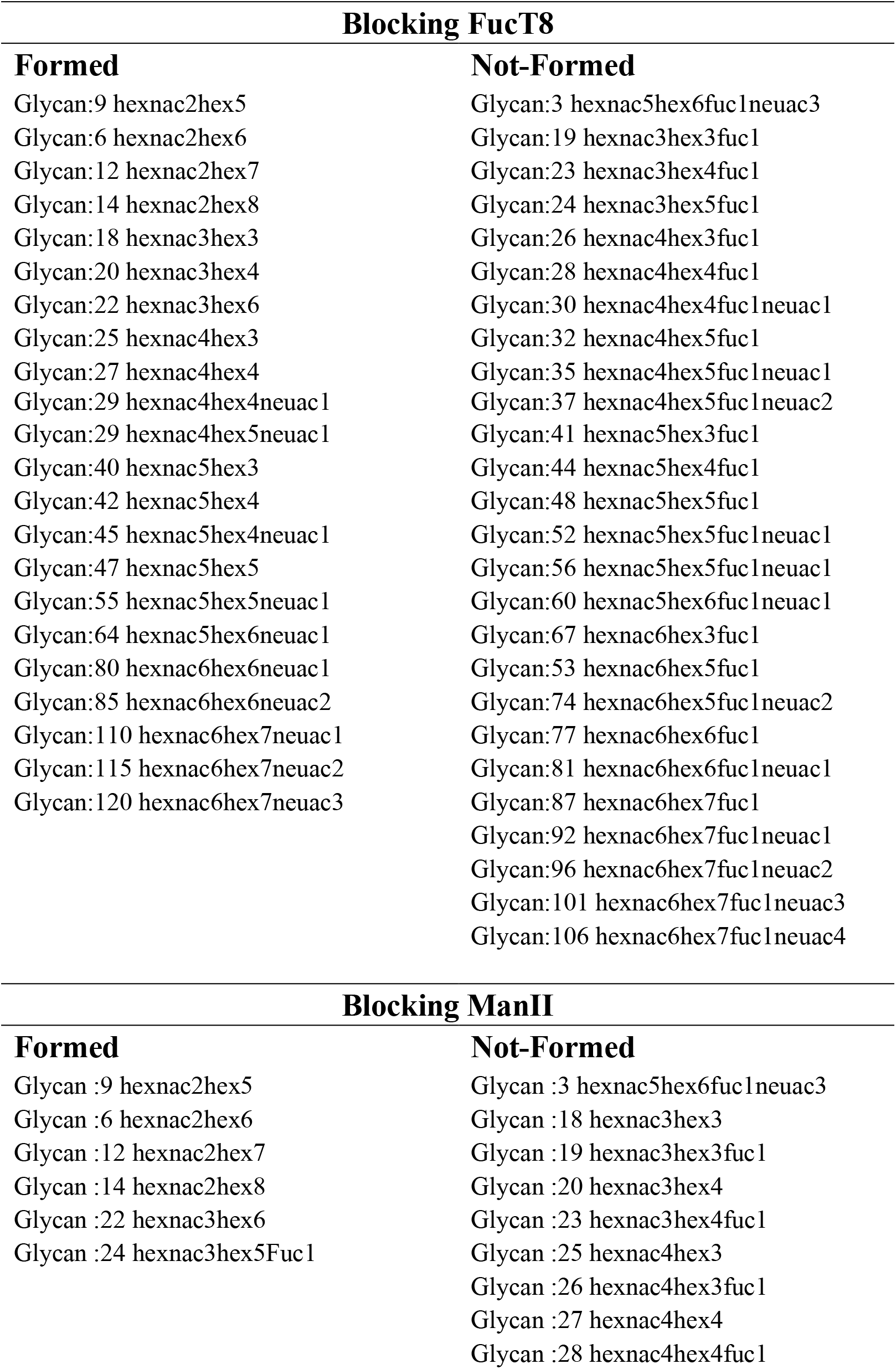
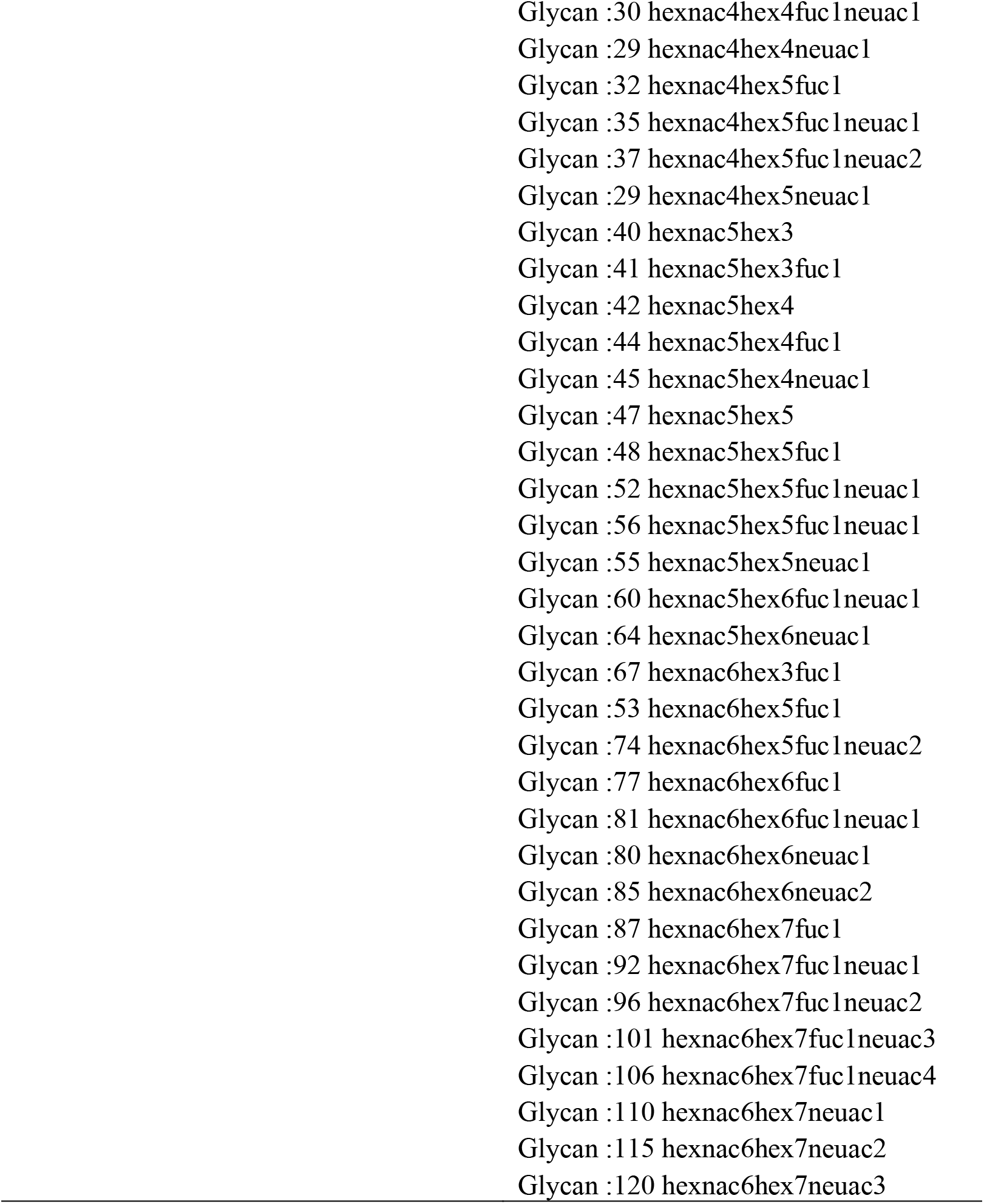
List of glycans that were formed vs not formed when specific enzymes were blocked when using data from Shajahan et al. Glycan numbers indicate nodes in Figure 2.

Even though we included data from both manuscripts, since we only used the representative glycan from Zhang et al, and a more complete version of data from Shajahan et al, it is likely that the latter is a more accurate representation of the glycosylation process of the virus.

## DISCUSSION

Here we present a preliminary N-glycan biosynthesis pathway of the spike S glycoprotein on SARS-CoV-2, which is not available currently for quick examination. We also share the individual glycan structures in a more easily accessible format for future glycoinformatics and molecular dynamics work. Based on our blocking simulation studies, the enzymes Man-II and FucT8 play important roles in the biosynthetic pathway, and without them the glycans synthesized are altered, changing the glycoprotein profile. The N-glycan biosynthesis pathway is a highly conserved two-step process – beginning in the endoplasmic reticulum, and ending in the golgi body ^22^. It has recently been recognized that the host glycosylation process is very relevant to vaccine development ^13^ and being able to identify the effect of modified glycans on the protein could further aid in development of vaccines. This first step will pave the way for such future work.

We did not choose to focus only on the glycans in the receptor binding domain (RBD) in the current report. Instead we showed that the overall glycan profile is altered dramatically by simulating glycosylation pathway inhibitors. Owing to the site-specific microheterogeneity that all glycoproteins express, which is also true of the SARS-CoV-2 spike protein, it becomes important in subsequent steps to study the effect of this variability on its pathogenicity, with specific focus on the glycans in the RBD. However, the role of these glycans in overall protein folding and dynamics is unclear, and still being studied. Several factors determine this microheterogeneity^23^, which in turn affects the structure, folding, and dynamics of the protein. So, it becomes important to not focus only on the glycans within the RBD of the spike S protein. Researchers have generated molecular dynamic simulations of the SARS-CoV-2 spike protein that represents site-specific micro-heterogeneity, based on mass spectrometry and imaging studies ^24,25^. Following up on our current approach, it would be possible to generate molecular dynamic simulation studies of altered spike S-glycoprotein that is generated by altering the glycosylation machinery, for vaccine development or other purposes.

Various levels of computational modeling of N-glycosylation have been used in the past. Here we used a pathway construction computational model. Alternatively, simulations that include the kinetics of enzyme activity by including rate equations can be more representative of the physiological process^26^. Using kinetics, it will further be possible to examine the effect of slowing down the rate of glycosylation of the spike S protein on overall viral replication^27^. Such an approach could shed light on potential pharmacological targets that would slow down both the host and the virus glycosylation pathways. If used appropriately, it may become possible to identify pharmacological targets that would affect the host the least, and the virus the most. Also, the time involved in testing several of these targets to identify ideal ones can be made short by using such network modeling tools before pre-clinical trials. However, this can be very challenging since the mammalian host post-translational glycosylation, and its downstream effect on proteins and their functions is still an active area of investigation.

Yet another approach is to use golgi-compartment representations within the modeling framework to model intra-organelle regional impacts on protein synthesis^28^. This could also aid in developing accurate representations of the enzymatic biosynthesis pathway. Alternately, for the current crisis, these approaches could evaluate competitive inhibitor glycans (natural or synthetic) and their effect on viral replication. By generating modified glycoproteins, it is possible to evaluate how they bind to, or alter the immune response of the host, since the host response to SARS-CoV-2 has been shown to be the determining factor in the severity of the manifested infection or in development of life-threatening adverse complications^29^. The added benefit of modeling is to be able to quickly narrow down targets by simulating several at once, while also knowing the underlying mechanism, which is not always possible in clinical studies. This makes computational modeling a useful tool in drug and vaccine discovery efforts.

## Limitations

As mentioned, there are several more approaches to construct N-glycosylation pathways. In addition to that, this work being computational, is preliminary, and requires further computational and basic/pre-clinical/clinical work to identify the effect of simulated outcomes. We did not conduct protein dynamic modeling studies, to determine if the altered glycan affects the protein, and its downstream binding with mammalian receptors.

## Acknowledgements

We would like to thank Prof. Sriram Neelamegham, University at Buffalo, for providing feedback on our manuscript, and Prof John Yin, University of Wisconsin-Madison for insight into virus replication processes. This work used the Extreme Science and Engineering Discovery Environment (XSEDE), which is supported by National Science Foundation grant number ACI-1548562.

